# Varying selection pressure for a Na^+^ sensing site in epithelial Na^+^ channel subunits reflect divergent roles in Na^+^ homeostasis

**DOI:** 10.1101/2024.02.16.580691

**Authors:** Xue-Ping Wang, Priyanka Srinivasan, Mustapha El Hamdaoui, Rafael Grytz, Ossama B. Kashlan

## Abstract

The epithelial Na^+^ channel (ENaC) emerged early in vertebrates and has played a role in Na^+^ and fluid homeostasis throughout vertebrate evolution. We previously showed that proteolytic activation of the channel evolved at the water to land transition of vertebrates. Sensitivity to extracellular Na^+^, known as Na^+^ self-inhibition, reduces ENaC function when Na^+^ concentrations are high and is a distinctive feature of the channel. A fourth ENaC subunit, δ, emerged in jawed fishes from an α subunit gene duplication. Here, we analyzed 846 α and δ subunit sequences and found that a key Asp in a postulated Na^+^ binding site was nearly always present in the α subunit, but frequently lost in the δ subunit (e.g., human). Analysis of site evolution and codon substitution rates provide evidence that the ancestral α subunit had the site and that purifying selection for the site relaxed in the δ subunit after its divergence from the α subunit, coinciding with a loss of δ subunit expression in renal tissues. We also provide evidence that the proposed Na^+^ binding site in the α subunit is a bona fide site by conferring novel function to channels comprising human δ subunits. Together, our findings provide evidence that ENaC Na^+^ self-inhibition improves fitness through its role in Na^+^ homeostasis in vertebrates.

## Introduction

Epithelial Na^+^ channels (ENaC) appeared in early vertebrates and play important roles in Na^+^ sensing and regulating extracellular fluids. The channel is typically expressed on cell surfaces, where its large extracellular domain can sense and respond to the extracellular environment (Kashlan et al., 2024). Accordingly, adaptive changes in ENaC function have paralleled vertebrate evolution. For example, cleavage sites that facilitate constitutive activation of the channel coevolved with the terrestrial migration of vertebrates, likely selected for by the increased threat of desiccation that terrestrial habitats pose (Wang et al., 2022). ENaC also directly senses Na^+^ and other small molecules present in extracellular fluids (Kashlan et al., 2024).

Inhibition by extracellular Na^+^, known as Na^+^ self-inhibition, was first observed in frog skin when rapid increases in extracellular Na^+^ initially increased currents as expected for a Na^+^ selective channel, but then reduced currents over a few seconds (Fuchs et al., 1977). Evidence supports Na^+^ binding at one or more extracellular effector sites that drive down the channel’s open probability (*P*_*o*_) (Kashlan et al., 2015; Sheng et al., 2006). In humans and mice, binding likely occurs at a site in the α subunit centered around human Asp-338 (mouse 365) in the β6-β7 loop of the β-ball domain (Figure 1). This site was first identified in mouse ENaC by examining the cation specificity of inhibition, effects on H^+^-dependent activation, and crosslinking experiments connecting local conformational changes to gating (Kashlan et al., 2015). Wichmann et al. confirmed key observations in both the α and δ subunits from *Xenopus laevis* (Wichmann et al., 2019). Noreng et al. observed electron density near human Asp-338 consistent with a bound cation and speculated that it was a Na^+^ ion (Noreng et al., 2020).

**Figure 1.**
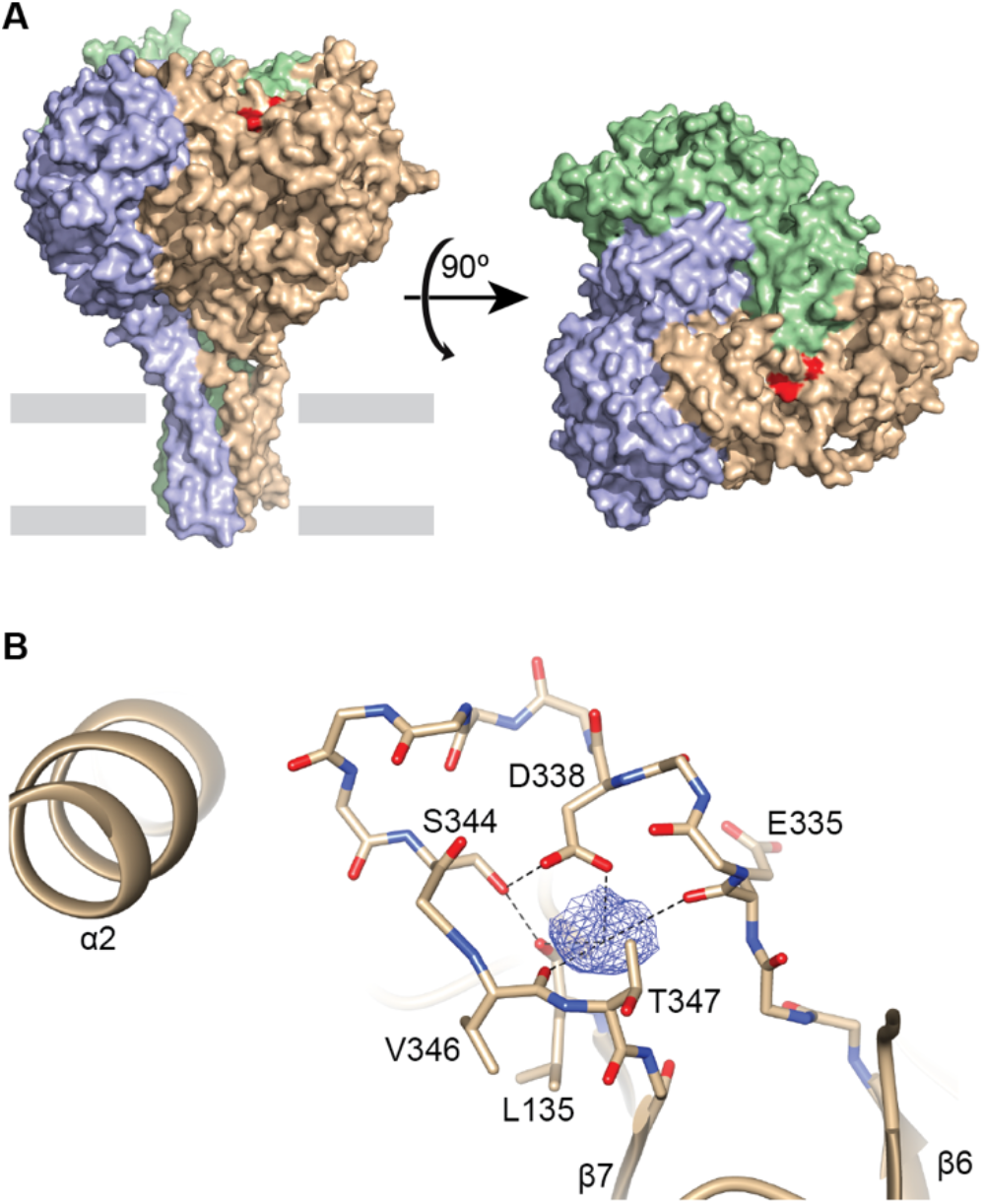
Cation binding site in the β6-β7 loop of the α subunit. (**A**) Surface model of human ENaC trimer (pdb code: 6BQN) made of α, β, and γ subunits, colored tan, green, and purple, respectively. Residues implicated in cation binding are colored red. Gray bars indicate approximate location of membrane. (**B**) Cartoon and sticks representation of cation binding pocket in the α subunit (pdb code: 6WTH). Carboxylate, hydroxide and carbonyl groups from residues in the β6-β7 loop and L135 contribute to the binding site. Dashed lines illustrate distances shorter than 3.5 Å.

ENaCs are heterotrimers composed from four paralogous subunits: α, β, γ, and δ. The α and δ subunits emerged from a gene duplication event in a jawed-fish ancestor (Wang et al., 2022). The δ subunit was subsequently lost in a subset of rodents including mice and rats (Gettings et al., 2021; Giraldez et al., 2012), which has led to a poor understanding of its physiological role. These two subunits are interchangeable in assembled channels and confer distinct functional properties. For example, human αβγ ENaC shows robust Na^+^ self-inhibition and is activated by proteases, whereas human δβγ ENaC is less sensitive to both Na^+^ and proteases (Haerteis et al., 2009; Ji et al., 2006). Human ENaC δ subunits lack an Asp at the analogous position associated with Na^+^ self-inhibition. Intriguingly, both the α and δ subunits from *Xenopus laevis*, rooted close to the α/δ subunit divergence point, retain the analogous Asp residue and demonstrate Na^+^ self-inhibition (Wichmann et al., 2019).

We hypothesized that residues involved in Na^+^ binding were present before the divergence of the α and δ subunits, and then selectively lost in the δ subunit after the divergence. Based on this, we hypothesized that a species closely related to humans retains the site and that we could resurrect a Na^+^ self-inhibition site in the human δ ENaC subunit. We determined the phylogenetic relationship of α and δ subunits and analyzed the appearance key residues at the proposed Na^+^ self-inhibition site. The central Asp demonstrated markedly different inheritance patterns after the divergence of the α and δ subunits. Analysis of the non-synonymous to synonymous substitution rate ratios (dN/dS) of the coding sequences provide evidence for purifying selection of residues comprising the site in the α subunit, but relaxed selection pressure for the analogous residues in the δ subunit. We identified a close human relative in the order *Scandentia* that retained the homologous Asp in its δ subunit and shows Na^+^ self-inhibition that depends on the same. After identifying site differences with human δ, we were able to confer Na^+^ self-inhibition to human δβγ channels through selective mutagenesis. We did not detect ENaC δ subunit expression in kidneys from the treeshrew, opossum or chicken, consistent with a loss of renal expression for the δ subunit after the divergence of α and δ subunits. Taken together, our data unambiguously define a Na^+^ self-inhibition site and provide evidence that Na^+^ self-inhibition improves fitness through its function in the kidney.

## Results

### Conservation of Asp associated with Na^+^-sensing

To examine the evolution of human αD338 and other residues implicated in Na^+^ binding for Na^+^ self-inhibition, we determined the phylogenetic relationship of more than 800 α and δ ENaC subunit sequences (Figure 2). We rooted the tree using the α subunit from the sea lamprey, which appeared before the divergence of α and δ subunits in a jawed fish (Wang et al., 2022). Our tree exhibits the expected phylogenetic relationships between the subunits based on this gene duplication event and speciation thereafter. When we examined the site aligning with human αD338 in the α subunits, we found an Asp in every sequence except in *Choloepus didactylus*. In the δ subunits, 59 species had a residue other than Asp at the same position. These include the Gymnophiona order of amphibians, all primates, several other mammalian groups, and several members of the Passerida suborder of perching birds. We also identified *Tupaia belangeri* (treeshrew) in the order Scandentia and *Galeopterus variegatus* (flying lemur) in the order Dermoptera as the closest human relatives with a δ subunit bearing an Asp in the same position in the alignment.

**Figure 2.**
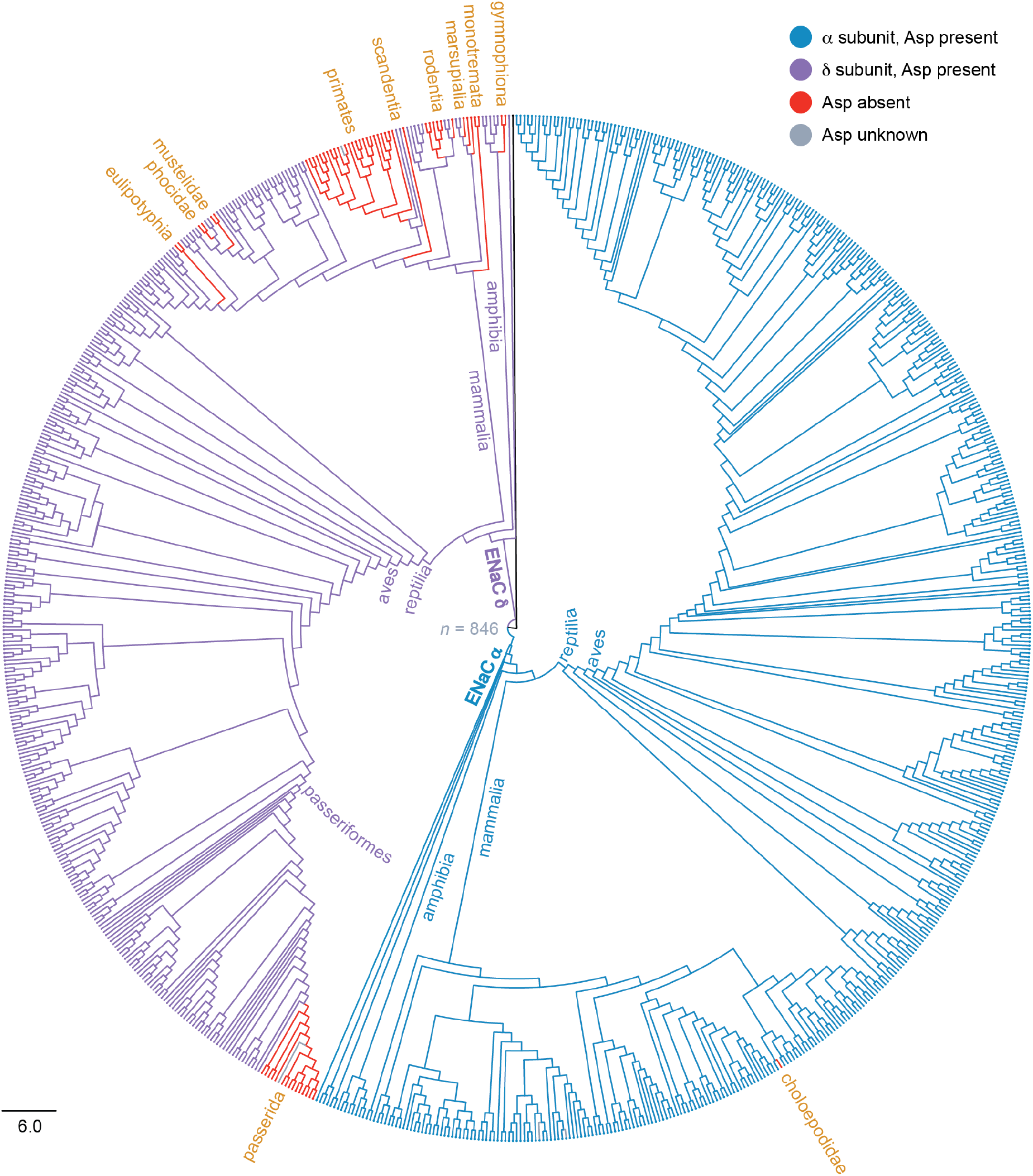
Phylogenetic analysis of ENaC α and δ subunits. Circular phylogram representation of maximum-likelihood tree calculated from 846 ENaC α and δ subunit protein sequences. The tree was rooted using the ENaC α subunit from sea lamprey. Branches are colored according to the residue aligning with human αAsp-338: Asp in α subunits are blue, Asp in δ subunits are purple, non-Asp residues are red, and unknown residues are gray. Key internal branch points and families are highlighted. Scale bar indicates the number of substitutions per site. **Supplementary file 1**. Accession numbers. **Supplementary file 2**. Tree file.

### Treeshrew δ exhibits Na^+^ self-inhibition

We hypothesized that the treeshrew δ subunit (tδ) has an intact Na^+^ binding site. We therefore synthesized and expressed tδ along with wild type human β (hβ) and γ (hγ) ENaC subunits in *Xenopus laevis* oocytes to assess its function using two-electrode voltage clamp current recordings. While clamping oocytes at -100 mV, we rapidly increased bath [Na^+^] from 1 mM Na^+^ to 110 mM Na^+^ in a 20 µL chamber perfused at 5 mL/min. This maneuver takes advantage of ENaC’s slow gating kinetics [∼0.2 opening events/s and 0.8 closing events/s (Anantharam et al., 2006)] relative to bath exchange rates (∼4 bath volumes/s) to measure Na^+^ self-inhibition (Figure 3A) (Chraibi & Horisberger, 2002). With 1 mM Na^+^ in the bath, ENaC’s *P*_*O*_ is relatively high. This results in an inward current peak when the driving force for inward currents is suddenly increased by an increase in bath [Na^+^]. Na^+^ then binds an extracellular effector site that drives down ENaC’s *P*_*O*_, rate limited by ENaC’s gating kinetics (Chraibi & Horisberger, 2002). Oocytes expressing tδ subunits exhibited robust Na^+^ self-inhibition (Figure 3B, C). To test whether the conserved Asp in tδ (tδD300) affects Na^+^ self-inhibition, we mutated it to its human δ equivalent, Pro. Mutation decreased the Na^+^ self-inhibition response, but did not abolish it (*p* = 0.007 for peak vs steady state currents by paired Student’s *t* test). These data are consistent with a role for tδD300 in Na^+^ binding, but also suggest residual Na^+^ binding in the tδD300P mutant.

**Figure 3.**
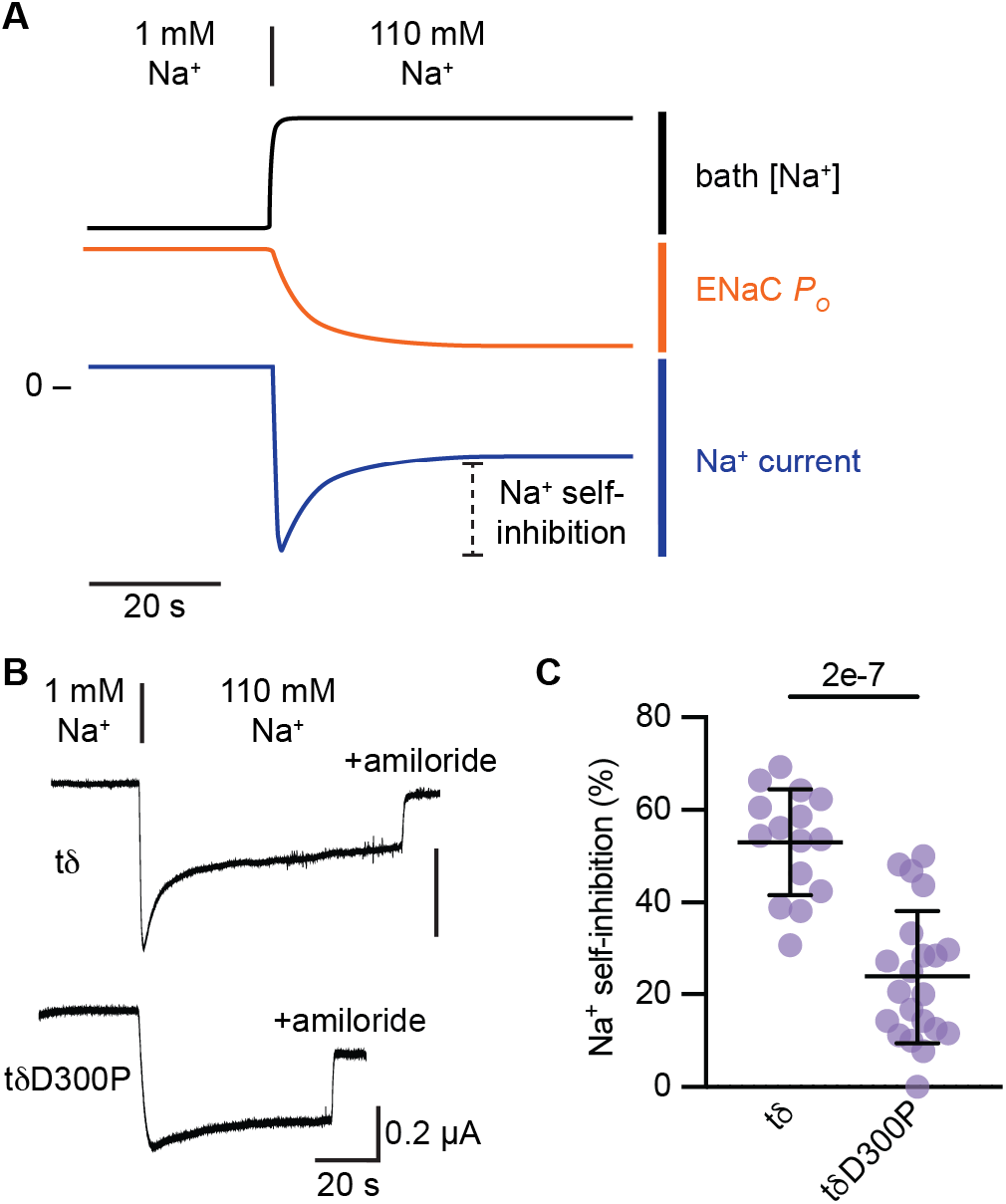
Conserved Asp in treeshrew δ EnaC subunit affects Na^+^ self-inhibition. (**A**) Schematic of voltage-clamp experiment to measure Na^+^ self-inhibition. With oocytes mounted in a 20 µL chamber perfused at 5 mL/min and clamped at -100 mV, bath [Na^+^] is rapidly increased from 1 mM Na^+^ to 110 mM Na^+^. This rapidly increases the driving force for Na^+^ entry and results in an inward current peak shortly thereafter. Simultaneously, extracellular Na^+^ binds ENaC and decreases *P*_*O*_ with time constant of ∼2-3 s. Declining currents from the peak over reflect decreases in *P*_*O*_. (**B**) Representative recordings of *Xenopus* oocytes injected with the tδ subunit indicated and complementary hβ and hγ EnaC subunits. 10 µM amiloride was added in 110 Mm Na^+^ buffer to determine ENaC-specific currents. (**C**) Na^+^ self-inhibition was assessed as the reduction of current from the peak to steady state 1 min after increasing bath [Na^+^] relative to the total amiloride-sensitive current measured from the peak to the value with amiloride. Groups were compared by Student’s *t* test.

### Selection pressure was lost for functional site residues in ENaC’s δ subunit

The pattern of Asp residues in our tree at the position aligning with human αD338 in extant species raises several questions. Do Asp gain loss rates differ between the α and δ subunits? Did ancestral subunits have an Asp at the site? What modes of selection are operating on residues implicated in Na^+^ binding?

To test for dependence of the site on subunit identity, we compared two nested models of site evolution over the full phylogenetic tree using BayesTraits and maximum likelihood methods (Pagel et al., 2004). We assigned each sequence as an α subunit or a δ subunit, and as either having or lacking an Asp according to Figure 2. For the 4 sequences in our tree lacking sequence information for the site (gray in Figure 2), we assigned ‘unknown’. The independent model, serving as the null hypothesis, contained three parameters: a site gain rate and site loss rate common to both subunits, and an appearance rate for the δ subunit. The dependent model added two free parameters by having independent rates for each subunit. A likelihood ratio test comparing the models strongly favored the dependent model (*p* = 0.02). In the favored model, the Asp gain rate:loss rate ratio in the α subunit was 68.7, favoring site retention, compared to 0.71 in the δ subunit, slightly favoring site loss.

To resolve whether the ancient ancestral subunit had an Asp at the site, we compared fits of the dependent model to the tree using MCMC methods while fixing the state of the site (present or absent) in the most recent common ancestor to the sea lamprey α, coelacanth α and coelacanth δ subunits. The parameter distribution shapes were similar for all three models (Figure 4, rows 1-3). However, fixing the ancient ancestor to lack the Asp increased the site gain rates, so that the rate distribution medians for the α and δ subunits were 25% and 35% greater, respectively, compared to the unconstrained model. Medians for the other parameters were within 8% of the unconstrained model. Fixing this early ancestor to have an Asp had little effect on model rates, with the medians for all rate distributions within 4% of the unconstrained model. We compared fits by converting marginal likelihoods from each MCMC run to log Bayes Factors, where values <2 provide weak evidence, >2 provide positive evidence, and >5 provide strong evidence of a preference between models. Fixing the ancient ancestor to have the Asp gave a log Bayes Factor of 0.7 vs the unconstrained model, while fixing the ancient ancestor to lack the Asp gave a log Bayes Factor of 4.7 vs the unconstrained model. Comparing the constrained models to each other gave a log Bayes Factor of 4.0, favoring the presence of an Asp in the ancestral node. These data provide evidence that the common ancestor at the base of the tree likely had an Asp at the site. We performed a similar analysis to determine whether the most recent common ancestor to human δ and treeshrew δ had an Asp at the site (Figure 4, rows 4-5). Comparing the model with the primate-treeshrew δ subunit ancestor constrained to lack the site to the unconstrained model (log Bayes Factor = 7.4) or to the model constrained to have the site in this ancestor (log Bayes Factor = 7.1) provides strong evidence that this more recent ancestor had the site, and that it was lost on the primate lineage after their divergence from Scandentia 73-100 million years ago (Zhou et al., 2015).

**Figure 4.**
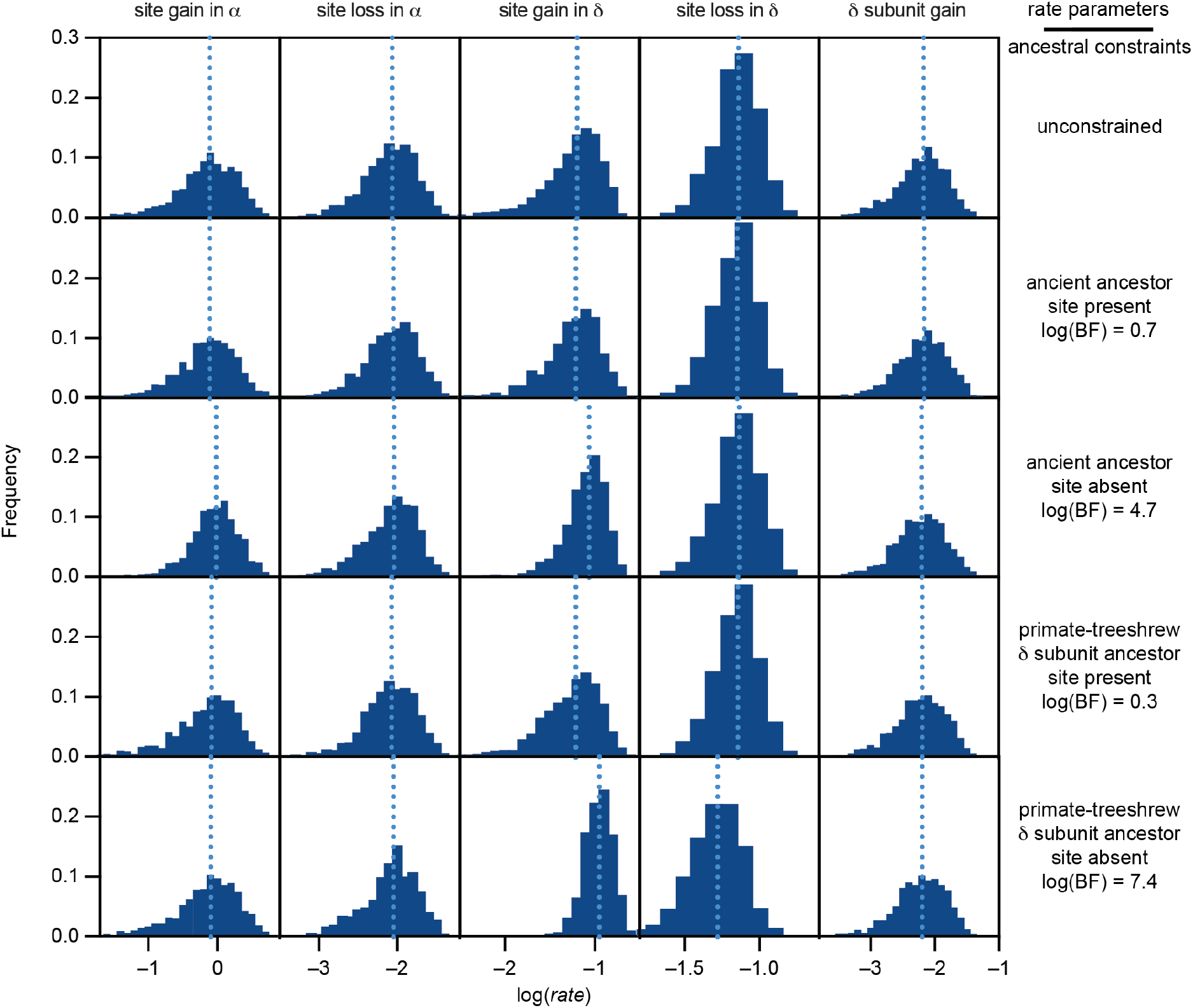
Fitted model parameters for Asp trait evolution. Trait evolution across the phylogenetic tree was determined using BayesTraits. MCMC runs were performed to determine the effect on model fits of fixing an ancient ancestor (the most recent common ancestor to sea lamprey α, coelacanth α, and coelacanth δ) or the primate-tree shrew δ subunit ancestor (the most recent common ancestor to human δ, treeshrew δ, and flying lemur δ) to either have or not have the Asp trait. Histograms of log-transformed rates from every 1,000^th^ iteration after the first 10,000 iterations are shown. The median values for each distribution are indicated with light blue dotted lines. Log Bayes Factors [log(BF)] are shown for each constrained model as compared to the unconstrained model. **Supplementary file 3**. Trait table. **Supplementary file 4**. Parameters for each BayesTraits run. **Supplementary file 5**. BayesTraits output files.

To determine whether a change in selection pressure after the divergence of the α and δ subunits explains the difference in site retention rates, we compared the non-synonymous substitution rate (*dN*) and synonymous substitution rate (*dS*) for the sequences encoding key residues implicated in Na^+^ self-inhibition (Figure 1). Values of *dN*/*dS* close to 1 are consistent with neutral drift, values greater than 1 indicate diversifying selection, and values less than one suggest purifying selection (Yang et al., 2000). We used HYPHY to measure and compare *dN* and *dS* for the key sites in each subunit (Kosakovsky Pond et al., 2020). Using HYPHY’s FEL module (Kosakovsky Pond & Frost, 2005), *dN*/*dS* was less than 1 for all selected sites, except for the δ subunit site aligning with hαV364 (Table 1). This is consistent with purifying selection at each of these sites. As the values for α appeared smaller than for δ, we then explicitly compared them using HYPHY’s Contrast-FEL module (Kosakovsky Pond et al., 2021). Values were smaller in the α subunit for sites aligning with hαL135, hαD338, hαS344, and hαV346. Strikingly, there was a 25-fold difference in *dN*/*dS* for the site aligning with hαD338. These data indicate that purifying selection pressure relaxed for several of the residues implicated in Na^+^ binding after the δ subunit diverged from the α subunit.

**Table 1.** Analysis of codon substitution rates. Analysis of *dN and dS* for residues implicated in Na^+^ binding (see Figure 1). *P*(*dN*<*dS*) was determined using FEL. The probability that *dN* for the α subunit is less than *dN* for the δ subunit, assuming a common *dS* [*P*(α < δ)], was determined using Contrast-FEL. **Supplementary file 6**. Nexus file for FEL and Contrast-FEL analysis. **Table 1–source data**. FEL and Contrast-FEL result files.

| Site (hα) | $\alpha$ subunit | | $\delta$ subunit | | $P(\alpha < \delta)$ |
| --- | --- | --- | --- | --- | --- |
| | $dN/dS$ | $P(dN < dS)$ | $dN/dS$ | $P(dN < dS)$ | |
| L135 | 0.015 | $< 0.001$ | 0.10 | $< 0.001$ | 0.03 |
| E335 | 0.062 | $< 0.001$ | 0.15 | $< 0.001$ | 0.1 |
| D338 | 0.020 | $< 0.001$ | 0.50 | 0.03 | $< 0.001$ |
| S344 | 0 | $< 0.001$ | 0.17 | $< 0.001$ | 0.002 |
| V346 | 0.12 | 0.004 | 0.74 | 0.39 | $< 0.001$ |
| T347 | 0.024 | $< 0.001$ | 0.060 | $< 0.001$ | 0.2 |

### Weakened selection pressure coincides with loss of renal δ ENaC subunit expression

We reasoned that ENaC Na^+^ self-inhibition is important for renal function since Na^+^ concentrations can vary widely in the distal nephron, e.g., 10-220 mM Na^+^ in tubular fluid from the rat’s distal nephron (Malnic et al., 1966). ENaC Na^+^ self-inhibition is well-tuned to this range, with apparent affinity weaker than 100 mM, and inhibitory effects evident at ∼10 mM Na^+^ (Chraibi & Horisberger, 2002; Sheng et al., 2006). Kidney lysates from *Xenopus laevis* frogs express both ENaC α and δ subunits (Wang et al., 2022). ENaC δ subunit expression in human kidneys appears to be low or absent (Menon et al., 2020). In treeshrew tissue lysates, we observed strong signals for the α, β, and γ subunit transcripts in kidney, but no signal for the δ subunit transcript (Figure 5A). Like human (Waldmann et al., 1995), we readily detected δ subunit transcript expression in treeshrew testis. Unlike human, we did not detect δ subunit transcripts in treeshrew brain, pancreas, or ovary tissues. We analyzed kidney tissue from *Marmosa mexicana*, a marsupial rooted close to the base of the mammals. As we were unable to amplify ENaC transcripts using primers based on related opossum species, we sequenced the transcripts in our kidney tissue sample. We identified transcripts for the α, β and γ ENaC subunits, but not the δ subunit, in *Marmosa mexicana* kidney tissue (Supplementary file 8; NCBI BioProject PRJNA1052681). As the δ subunit from several birds also lack the key Asp, we analyzed tissues from *Gallus gallus* (Figure 5B). Similar to the treeshrew, we detected transcripts for the α, β, and γ subunits, but not the δ subunit, in chicken kidney lysates. We did detect a faint signal for the δ subunit in chicken colon and spleen tissues, where we also detected signals for the α and γ subunit, but not the β subunit. As we observed δ subunit expression in kidneys from frog, but not human, treeshrew, opossum, or chicken, our data suggest that renal δ subunit expression was lost in reptiles, birds and mammals after their divergence from amphibians.

**Figure 5.**
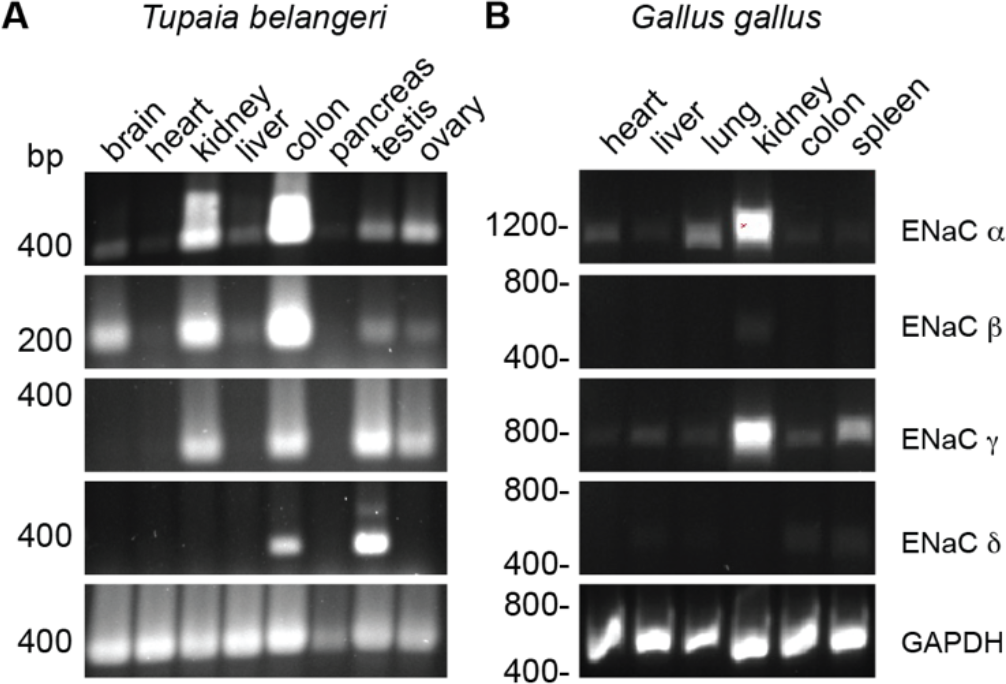
Tissue distribution of ENaC subunit transcripts in *Tupaia belangeri* (treeshrew) and *Gallus gallus* (chicken). Tissue homogenates were used to generate cDNA libraries. PCR reactions were performed using primer pairs indicated in Supplementary file 7. **Supplementary file 7**. Primers for RT-PCR and expected product sizes. **Figure 5–source data**. Uncropped gels.

### Na^+^ binding site resurrection in human δ ENaC

Given the likely presence of Na^+^ self-inhibition in the most recent common ancestor to tδ and hδ, and its loss in primates thereafter, we hypothesized that we could reconstruct a Na^+^ binding site in hδ and confer Na^+^ self-inhibition to channels comprising hδ subunits. We identified key differences in the cation binding pocket between the hδ and tδ subunits at the site, and mutated hδ accordingly (Figure 6A). Mutating hδP314 to Asp produced a discernable current peak after increasing extracellular [Na^+^] (Figures 6B, C). Additionally mutating hδG136 to Met and/or hδL322 to Glu further increased Na^+^ self-inhibition, so that responses for channels comprising double and triple hδ subunit mutants were similar to channels comprising the hα subunit. Each hδ mutant increased the response to extracellular Na^+^ compared to wild type hδ. These data conferring novel function to hδ provide evidence that hαD338 and neighboring residues (Figure 1) constitute a bona fide inhibitory Na^+^ effector site.

**Figure 6.**
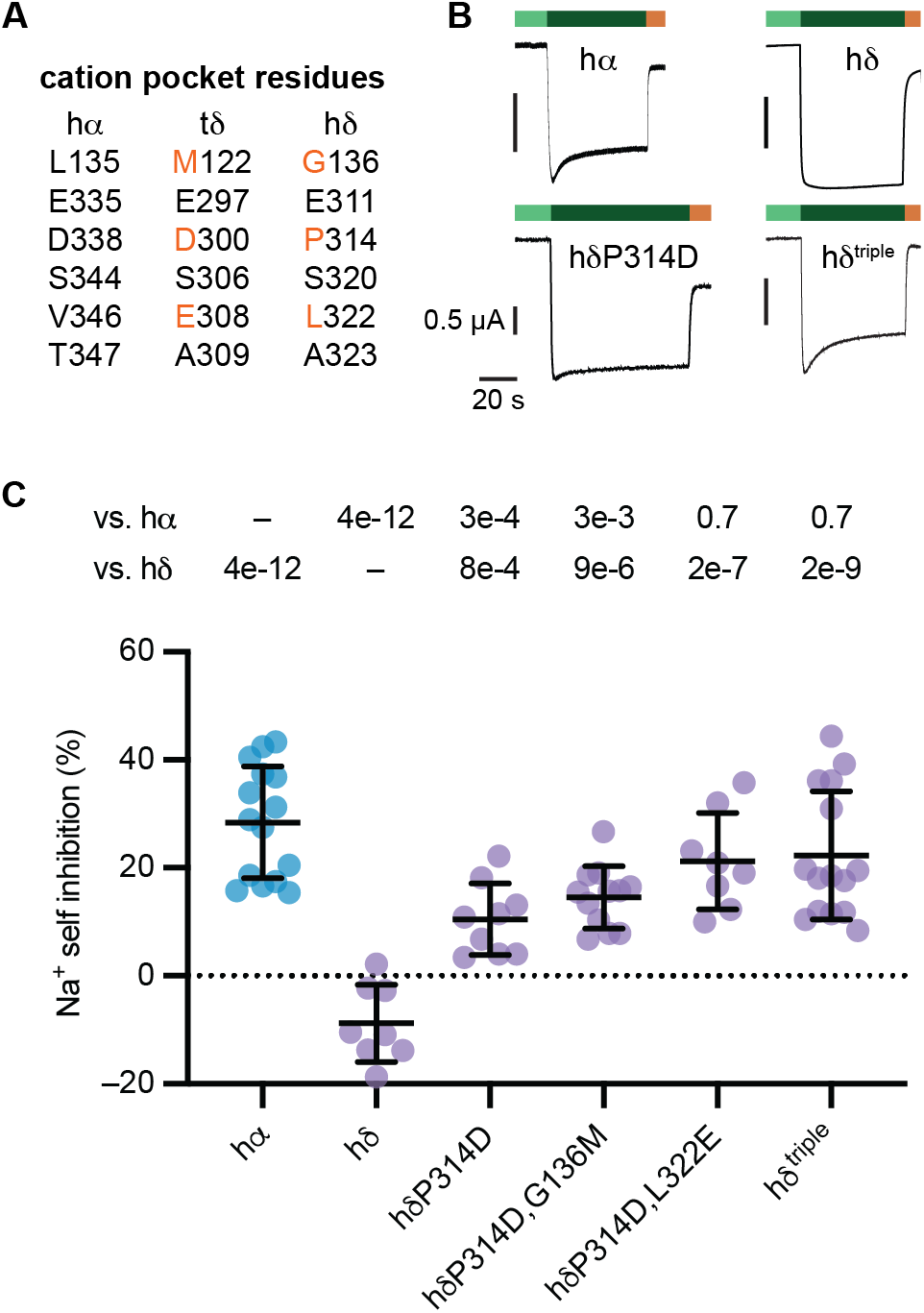
Na^+^ binding site resurrection in the δ subunit of human ENaC. (**A**) Key resides involved in the cation binding pocket in hα, along with their tδ and hδ equivalents. (**B**) Representative recordings of *Xenopus* oocytes injected with the hα or hδ subunit indicated and complementary hβ and hγ ENaC subunits. Bath conditions are indicated using colored bars: 1 mM Na^+^ (light green), 110 mM Na^+^ (dark green), and 10 µM amiloride in 110 mM Na^+^ buffer (orange). (**C**) Na^+^ self-inhibition was assessed as the relative loss of amiloride-sensitive current from the peak to steady state, as above, and groups were compared using one-way ANOVA and Šidáks multiple comparisons test.

## Discussion

ENaC subunits emerged early in vertebrates, with α, β, and γ subunits present in jawless fishes (Figure 7). Our findings provide evidence that this ancestral ENaC was inhibited by Na^+^ binding to an extracellular site in the channel’s α subunit, and that this functional property improved fitness throughout vertebrate evolution. Although first observed five decades ago (Fuchs et al., 1977), the physiological relevance of ENaC Na^+^ self-inhibition remains poorly understood (Kashlan et al., 2024). Its relevance has been linked to ENaC’s sensitivity to other regulatory factors, e.g., proteases, activation through which has been proposed to depend on Na^+^ self-inhibition (Noreng et al., 2020; Sheng et al., 2006). However, the apparent functional link can also be explained through indirect mechanisms (Kashlan et al., 2010; Passero et al., 2010).

**Figure 7.**
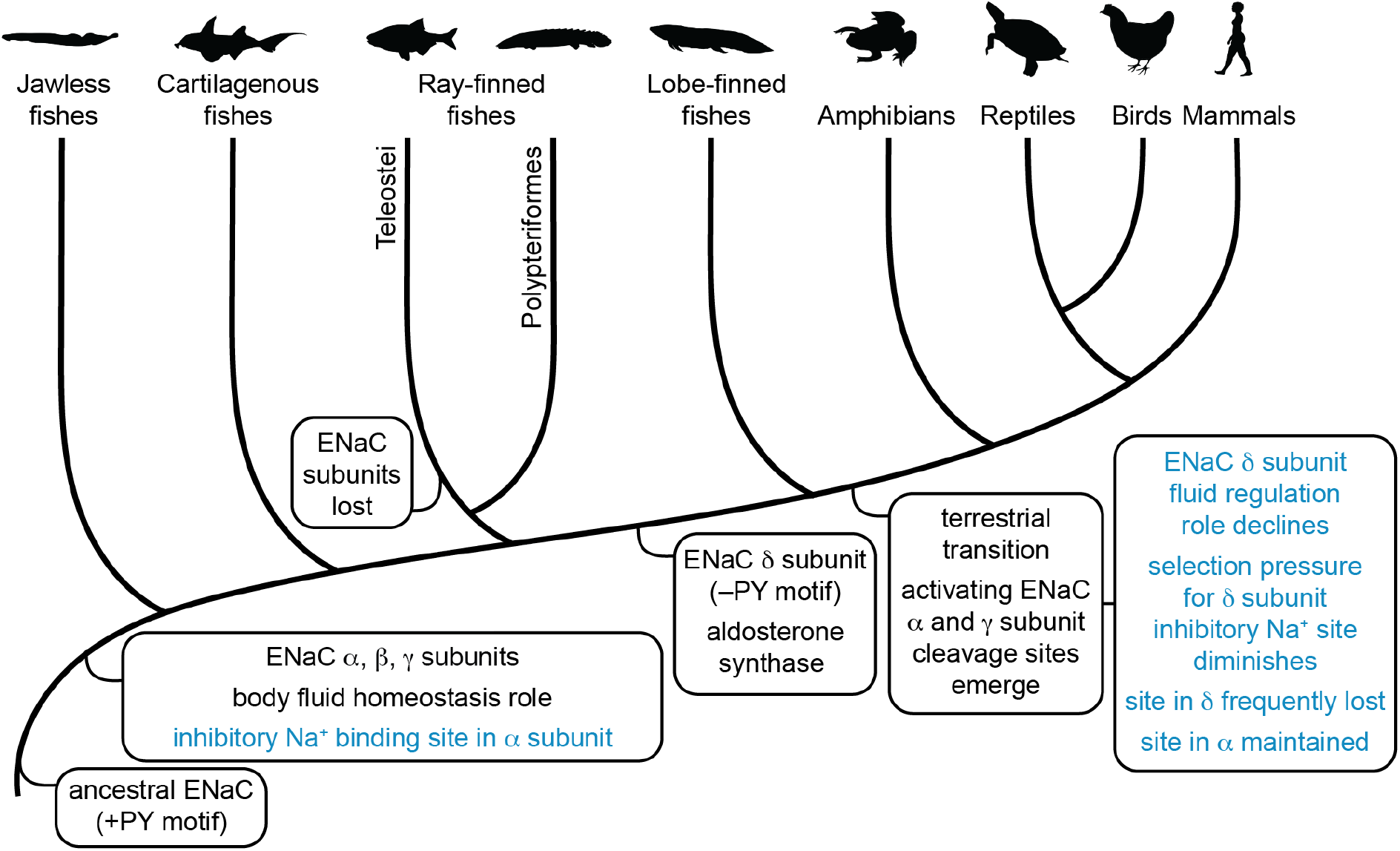
Evolution of ENaC subunits and regulatory motifs. Finding in the current work are highlighted in blue. Distinct ENaC α, β, and γ subunits appeared early in vertebrate evolution, with expression consistently found in organs responsible for maintaining Na^+^ homeostasis. Each of these subunits possessed a C-terminal PY motif that facilitated regulation of channel abundance by Nedd4-2. We found that the ancestral channel also likely had an inhibitory Na^+^ binding site in its α subunit that made the channel sensitive to extracellular Na^+^. ENaC subunits were lost in ray-finned fishes after the divergence of bichirs. In a jawed fish ancestor, the δ subunit emerged from an α subunit gene duplication event. As extant δ subunits lack a PY motif, the ancestral δ subunit likely lost its PY motif shortly after first emerging. Aldosterone synthase, required for aldosterone signaling also appears at this time. Aldosterone regulates ENaC through several mechanisms, including Nedd4-2, which predates aldosterone, and activating ENaC cleavage in the α and γ subunits, which emerged afterwards. Motifs for both are absent in the δ subunit. We also found that many δ subunits lack the Na^+^ binding motif required for Na^+^ self-inhibition. This was due to weakened selection pressure to maintain the Na^+^ binding site in the δ subunit, as compared to the α subunit. This loss coincides with a loss of δ subunit expression in kidney tissues, while α subunit expression was maintained. These observations suggest that ENaC’s role in total body Na^+^ and fluid homeostasis underlie the selection pressure that maintained the Na^+^ binding site in the α subunit. Figure adapted from previous work (Wang et al., 2022). Animal silhouettes courtesy of PhyloPic (http://www.phylopic.org).

Furthermore, the evolution of ENaC functional sites does not support this (Figure 7). Purifying selection maintained an inhibitory Na^+^ binding site in the α subunit throughout vertebrate evolution, both before and after sites for proteolytic activation emerged in the α and γ subunits with the migration of vertebrates to terrestrial habitats (Wang et al., 2022). Unlike proteolytic activation, proton-dependent activation is directly connected to Na^+^ self-inhibition, since protons activate the channel by directly competing with Na^+^ for the key Asp (Collier & Snyder, 2009; Kashlan et al., 2015; Wichmann et al., 2019). However, ENaC pH-sensitivity varies widely between species and protonation of the Asp involved in Na^+^ binding appears to be only part of the mechanism (Collier et al., 2012; Wichmann et al., 2019). We therefore conclude that ENaC sensitivity to extracellular Na^+^ *per se* improves fitness.

We propose that the loss of renal expression caused the relaxation of purifying selection for Na^+^ binding residues in the δ subunit. After the divergence of the α and δ subunits, δ subunit expression in renal tissues appears to have been lost. We detected renal δ subunit expression in *Xenopus laevis* frogs, while we and others detected little or no δ subunit expression in human, treeshrew, opossum or chicken kidney tissues (Figure 5) (Giraldez et al., 2007; Menon et al., 2020; Wang et al., 2022). Conversely, the α subunit is expressed in the kidneys of each of these species (Menon et al., 2020; Wang et al., 2022), and in the kidneys of the ropefish and two species of lungfish, which do not have ENaC δ subunits (Uchiyama et al., 2015; Uchiyama et al., 2012; Wang et al., 2022). The notion that ENaC Na^+^ self-inhibition improves renal function is attractive. Na^+^ concentrations in the distal renal tubule are highly variable, unlike many other sites that express ENaC where Na^+^ concentrations are constitutively high, such as in lung fluid and blood. Tonic inhibition through Na^+^ self-inhibition could increase fitness under these conditions, though a modest loss-of-function mutation could produce a similar effect. On the other hand, the apparent affinity for Na^+^ binding to ENaC, while weak, is well-matched to the distal renal tubule Na^+^ concentrations measured in rats (Malnic et al., 1966). The net effect of Na^+^ self-inhibition is to maximize Na^+^ transport when extracellular Na^+^ concentrations are low, and dampen Na^+^ transport when extracellular Na^+^ concentrations are high. This effectively sharpens the channel’s Na^+^ saturation curve so that Na^+^ transport rates rise steeply as Na^+^ concentrations increase to 20 mM, but then plateau at higher concentrations (Chraibi & Horisberger, 2002; Sheng et al., 2006). A Na^+^ channel tuned to efficiently recover Na^+^ when Na^+^ is scarce, while limiting recovery when Na^+^ is abundant may have provided a selective advantage. Further experimentation will be required to determine the role of Na^+^ self-inhibition in renal physiology.

Na^+^ self-inhibition appears to define ancestral ENaCs. These early channels also had asymmetric ion conduction pathways conferring high Na^+^:K^+^ selectivity, and C-terminal PPxY motifs that facilitate interactions with NEDD4-2 (Hanukoglu & Hanukoglu, 2016; Wang et al., 2022; Wesch et al., 2010). Aldosterone signaling later co-opted NEDD4-2 dependent ENaC regulation (Wang et al., 2022). In ENaC expressing fishes, the channel mediates Na^+^ uptake for body fluid homeostasis, and accordingly, its subunits have been found in gill, kidney and intestinal tissues (Ferreira-Martins et al., 2016; Tseng et al., 2022; Uchiyama et al., 2015; Uchiyama et al., 2012; Wang et al., 2022). Ray-finned fishes lost ENaC subunits after the divergence of bichirs (Hwang et al., 2011)(Wang et al., 2022), possibly because Na^+^ uptake through Na^+^-H^+^ exchanger 3 (NHE3) was more energetically efficient (Tseng et al., 2022). When δ subunits emerged in jawed fishes, the new protein quickly lost its C-terminal PPxY motifs (Wang et al., 2022; Wesch et al., 2010) and later lost Na^+^ self-inhibition. Its tissue distribution also changed. In humans, δ subunit expression is low or absent in aldosterone-sensitive tissues (e.g., kidney and colon), and abundant in the central nervous system (Giraldez et al., 2007; Kashlan et al., 2024; Waldmann et al., 1995; Yamamura et al., 2006). Disparate resting membrane potential requirements in transporting epithelia as compared to excitable cells may alternatively account for the differences in selection we found. ENaC expression increases membrane Na^+^ permeability and promotes depolarization, with Na^+^ self-inhibition limiting the effect at higher Na^+^ concentrations. ENaC-dependent membrane depolarization is important in several contexts, including in the distal nephron, and excitable cells in the lingual epithelium and central nervous system (Giraldez et al., 2013; Gray et al., 2005; Nomura et al., 2020; Oshima et al., 2013). However, in the treeshrew we detected δ subunit expression in the colon, but not the brain or kidney. Furthermore, the δ subunit was lost altogether in several rodent species (Gettings et al., 2021). Finally, we observed a loss of purifying selection for Na^+^ self-inhibition in the δ subunit but did not find evidence for diversifying selection. We therefore find no evidence that the loss of Na^+^ self-inhibition in the δ subunit increased fitness. Instead, evidence suggests that the loss of Na^+^ self-inhibition in the δ subunit had little impact, in contrast to the α subunit. If the δ subunit has an evolutionarily conserved role in physiology, it remains undefined.

In summary, we conferred novel function to the ENaC δ subunit, providing evidence that the proposed Na^+^ binding site in the α subunit’s periphery is a bona fide allosteric inhibitory effector site. Purifying selection pressure maintained this site in the α subunit but relaxed in the paralogous δ subunit leading to site loss on numerous occasions. This site was likely present in ancient ENaCs when vertebrates first emerged and may be critical to the channel’s role in Na^+^ homeostasis.

## Materials and Methods

### Multiple sequence alignment and phylogenetic tree calculation

Sequences were retrieved using NCBI’s BLAST tool (https://blast.ncbi.nlm.nih.gov/Blast.cgi) and human ENaC α and δ subunit protein sequences (NCBI accession: NP_001029.1, AAI25075.1), resulting in 846 protein sequences (Supplementary file 1). Sequences were then aligned using Mafft (Katoh & Standley, 2013), and curated using BMGE (Criscuolo & Gribaldo, 2010). The phylogenetic tree was calculated from the curated alignment using PhyML (Guindon et al., 2010) and the LG substitution model (Le & Gascuel, 2008). The resulting tree (Supplementary file 2) was visualized using FigTree (http://tree.bio.ed.ac.uk/software/figtree/) and used in subsequent analyses.

### Testing phylogenetic models of site gain and loss

Evolutionary models of two binary traits were compared using BayesTraits V3 (www.evolution.rdg.ac.uk) (Pagel et al., 2004). Traits were assigned as either α or δ subunits, and as having or lacking the Asp, as indicated in Figure 2 (Supplementary file 3). Sequences where the key region was missing (gray in Figure 2) were assigned as unknown. Sea lamprey α (black in Figure 2) was assigned as an α subunit having the Asp trait. Parameters for each BayesTraits run are provided in Supplementary file 4 and results are provided in Supplementary file 5. We compared nested models using a likelihood ratio test, with 2 degrees of freedom due to the 2 extra parameters in the dependent model. The likelihood ratio statistic [2 × (log-likelihood⟨dependent model⟩ − log-likelihood⟨independent model⟩)] was converted to a *p* value using Microsoft Excel and the *chisq*.*dist*.*rt* function.

To determine whether the Asp trait was present in an ancestral node, we used BayesTraits to fit the dependent model using MCMC methods while fixing the node of interest to reflect each hypothesis. Parameters for each run are provided in Supplementary file 4. Likely average values for model parameters were determined using maximum likelihood runs for each model. Exponentially distributed priors with a mean rounded to the nearest order of magnitude of the predicted mean were used for all parameters. Runs were performed for the default number of iterations (1,010,000), and resulted in parameter distributions with medians similar to preliminary maximum likelihood runs, consistent with convergence. Log marginal likelihood values were calculated using the stepping stone sampler (100 stones with 10,000 iterations). Evidence for model preference was determined using log Bayes Factors, calculated as: 2 × (log marginal likelihood model 1 − log marginal likelihood model 2), where values <2 provide weak evidence, values >2 provide positive evidence, and values >5 provide strong evidence for a model preference. *dN/dS measurements*

Values for *dN* and *dS* were measured and compared using HYPHY’s FEL and Contrast-FEL modules implemented at datamonky.org. Codon sequences were retrieved for 458 α and δ subunit sequences, and then aligned and trimmed to the region of interest (Supplementary file 6). The phylogenetic relationship of the sequence subset was determined from the protein sequences, as described above, and appended to the sequence file. After uploading sequence and phylogenetic data, the user defined tree was rerooted on the lamprey α sequence. To compare dN and dS for each subunit, the α subunit or δ subunit branches were selected before running FEL. To compare *dN* between subunits, α subunit and δ subunit partitions were defined before running Contrast-FEL. Results for FEL and Contrast-FEL runs are provided in Table 1– Source data, with summary data provided in Table 1.

### Site-directed mutagenesis and ENaC expression in Xenopus oocytes

Plasmids encoding wild type human ENaC subunits (hα, hβ, hγ, and hδ) were gifts from Tom Kleyman. A plasmid encoding tδ was synthesized in pTwist CMV plasmid (Twist Bioscience) and moved into psp64 with a C-terminal HA epitope tag. Point mutations were generated using the QuikChange II XL site-directed mutagenesis kit (Agilent, Santa Clara, CA) and confirmed by direct sequencing (GENEWIZ). Wildtype and mutant RNAs were synthesized using mMESSAGE mMACHINE T3, T7 or Sp6 *in vitro* transcriptional kit (Invitrogen). Oocytes from *Xenopus laevis* frogs were harvested as previously described (Sheng et al., 2005), as approved by the University of Pittsburgh’s Institutional Animal Care and Use Committee and were provided by the Pittsburgh Center for Kidney Research. 2 ng of each ENaC wildtype or mutant subunit was injected into stage V or stage VI oocytes maintained at 18 °C in modified Barth’s saline solution: 88 mM NaCl, 1 mM KCl, 2.4 mM NaHCO_3_, 15 mM Hepes, 0.3 mM Ca(NO_3_)_2_, 0.41 mM CaCl_2_, 0.82 mM MgSO_4_, 10 μg/mL streptomycin sulfate, 100 μg/mL gentamycin sulfate, and 10 μg/mL sodium penicillin, pH 7.4.

### Measurement of Na^+^ self-inhibition by two-electrode voltage clamp

Two-electrode voltage clamp studies were performed at room temperature 24 h after RNA injection using an Axoclamp 900A Computer-Controlled Microelectrode Amplifier and DigiData 1440A interface controlled by pClamp 10.4 (Molecular Devices). Oocytes were placed in a chamber and perfused at constant flow rates (5 mL/min). Glass pipettes filled with 3 M KCl were inserted into oocytes, and the intracellular potential was clamped at −100 mV. Na^+^ self-inhibition measurements were performed as previously described (Kashlan et al., 2018). Briefly, Na^+^ self-inhibition responses were recorded following a rapid transition from a low [Na^+^] bath solution (1 mM NaCl, 109 mM *N*-methyl-d-glucamine, 2 mM KCl, 2 mM CaCl_2_, and 10 mM HEPES, pH 7.4) to a high [Na^+^] bath solution (110 mM NaCl, 2 mM KCl, 2 mM CaCl_2_, and 10 mm HEPES, pH 7.4). Towards the end of each recording, amiloride (10 µM) was added to define the ENaC-dependent portion of the current. The currents at the peak and at steady-state were used to determine the Na^+^ self-inhibition response. Individual data points and shown with summary statistics (mean ± SD). Significance between two groups was determined by Student’s t test and for multiple groups by one-way ANOVA followed by Šidáks multiple comparisons test. *P*<0.05 was considered significant. All statistical analyses were performed using GraphPad Prism 10.

### Tissue distribution of ENaC subunit transcripts

Northern treeshrew (*Tupaia belangeri*) tissues were harvested from ∼30-week-old animals bred through the University of Alabama at Birmingham (UAB) treeshrew core, as approved by the UAB Institutional Animal Care and Use Committee. *Gallus gallus* tissues were harvested from a female slaughtered at Double R Ranch (New Wilmington, Pennsylvania). *Marmosa mexicana* kidney tissue was acquired through the Carnegie Museum of Natural History, which was collected from an adult female specimen in Izabel, Guatemala at 15º40’N, 88º41’W on June 13, 1994, and stored at -80ºC.

Total RNA was extracted from approximately 50 mg tissue using TRIzol reagent (Invitrogen). cDNA was generated using the RevertAid Reverse Transcriptase Kit (ThermoFisher Scientific). Specific primers for ENaC subunits, and glyceraldehyde-3-phosphate dehydrogenase (GAPDH) as an internal control (Supplementary file 7) were designed using NCBI primer-blast tools. PCR was run for 30 cycles using GoTaq G2 Green master mix (Promega) and specific primers, with the annealing temperature set at 58°C. PCR products were visualized with GelRed dye (Biotium) after agarose gel electrophoresis using a GelDoc imaging system (BioRad).

## Abbreviations

ENaC: epithelial Na^+^ channel,
*P*_*O*_: open probability,
*dN*: nonsynonymous substitution rate,
*dS*: synonymous substitution rate,
ENaC: subunit, tδ; treeshrew δ
hα, hβ, hγ,: or hδ human ENaC subunits:

## Supplementary files

**Supplementary file 1**. Accession numbers.

**Supplementary file 2**. Tree file.

**Supplementary file 3**. Trait table.

**Supplementary file 4**. Parameters for each BayesTraits run.

**Supplementary file 5**. BayesTraits output files.

**Supplementary file 6**. Nexus file for FEL and Contrast-FEL analysis.

**Supplementary file 7**. Primers for RT-PCR and expected product sizes.

**Supplementary file 8**. Alignment of *Marmosa mexicana* sequences (NCBI BioProject PRJNA1052681) with *Homo sapiens* ENaC subunits [NCBI accession numbers NP_001029.1 (ENaC α), NP_000327.2 (ENaC β), NP_001030.2 (ENaC γ)]. No sequences for the δ subunit were identified from *Marmosa mexicana* kidney tissue.

## Source data

**Figure 5–source data**. Uncropped gels.

**Table 1–source data**. FEL and Contrast-FEL result files.

## Data availability

Data for electrophysiology and uncropped gels are provided as source data. Primer sequences are provided as a supplementary file. Sequencing data for *Marmosa mexicana* kidney tissue are available at NCBI (BioProject PRJNA1052681). Input files and parameters used for phylogenetic trait analysis are provided as supplementary files.

## Acknowledgements

We thank Suzanne McLaren at the Carnegie Museum of Natural History for providing *Marmosa mexicana* tissue, and Anne Ruxandra-Carvunis, Allyson O’Donnell, Catherine Baty, Arohan Subramanya, Thomas Kleyman, and Evan Ray for helpful suggestions.

