## Supplemental Data 1 for "Varying selection pressure for a Na^+^ sensing site in epithelial Na^+^ channel subunits reflect divergent roles in Na^+^ homeostasis"

*Tupaia belangeri* RT-PCR

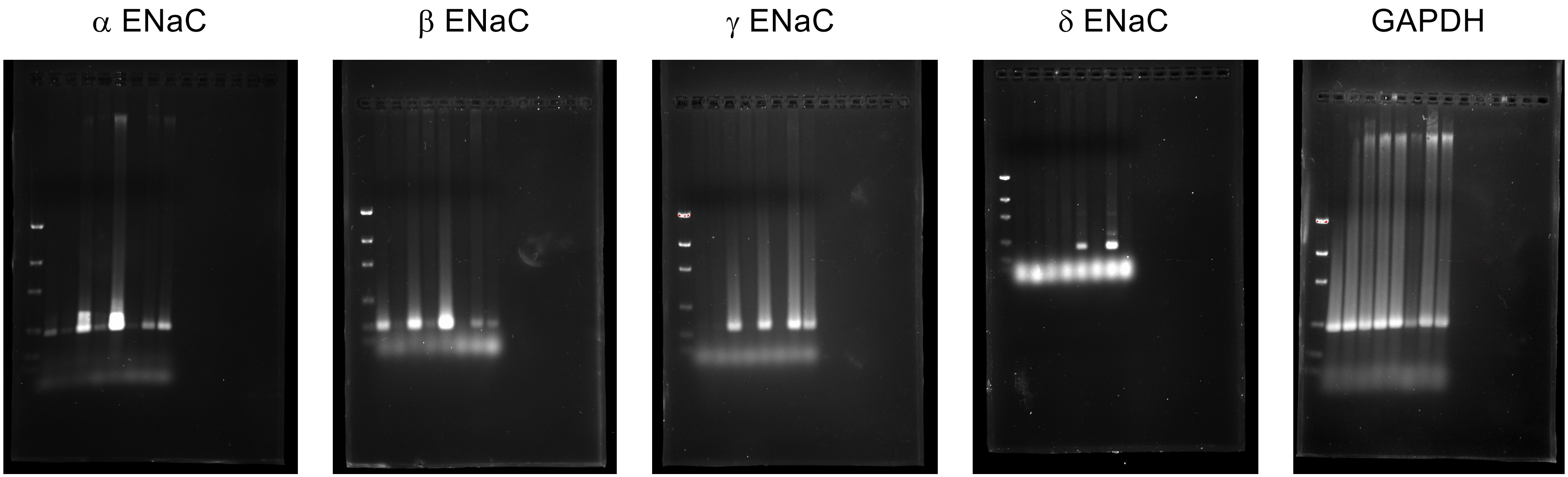


Markers: 2k, 1.2k, 800, 400, 200, 100
Lane order: markers, brain, heart, kidney, liver, colon, pancreas, testis, ovary

*Gallus gallus* RT-PCR

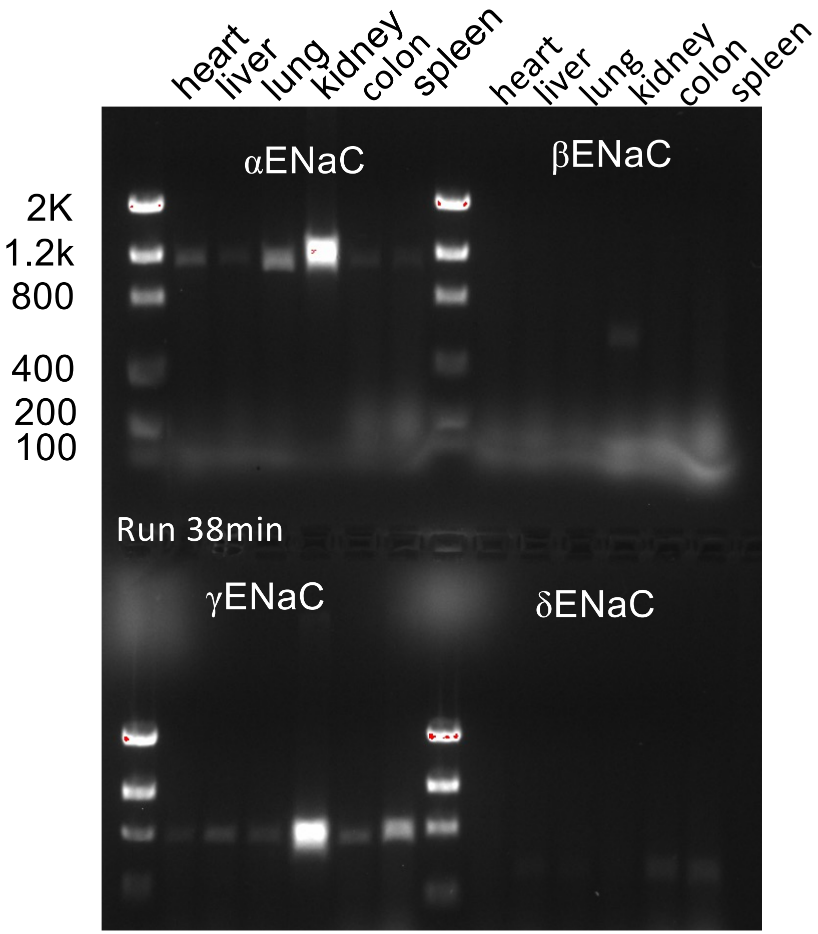


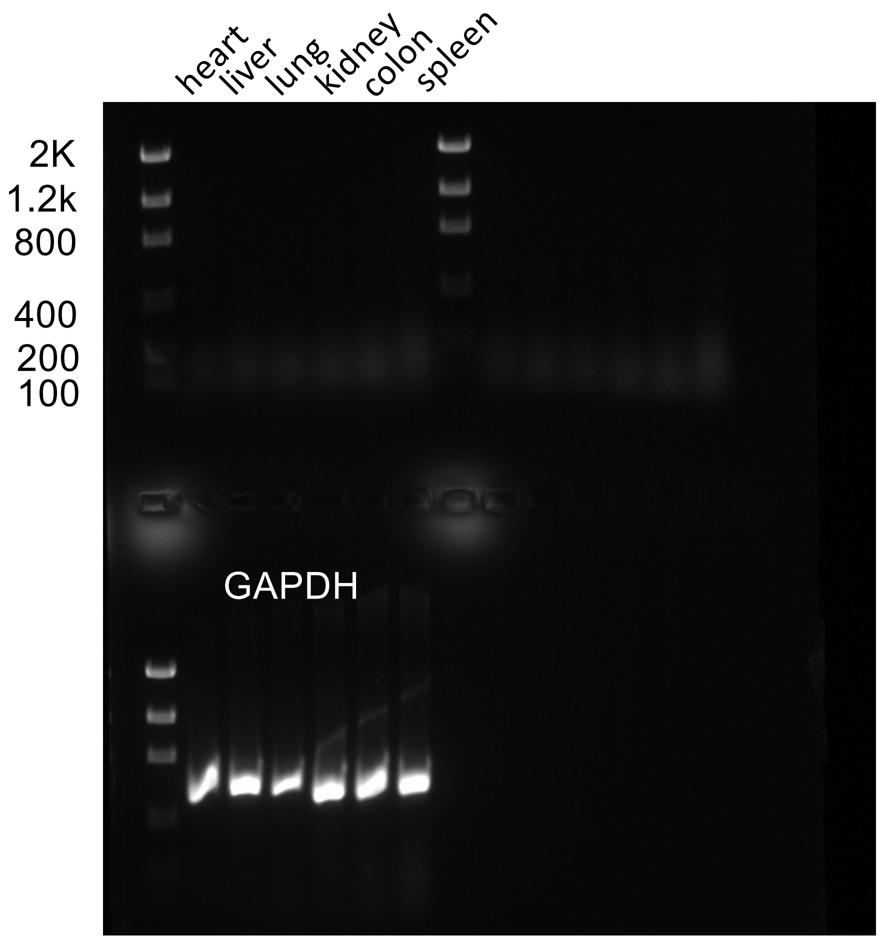
