## Supplementary file 7 for "Varying selection pressure for a Na^+^ sensing site in epithelial Na^+^ channel subunits reflect divergent roles in Na^+^ homeostasis"

Primers for RT-PCR and expected product sizes.

| Target | Primer sequence (5'–3') | | Amplicon (bp) |
| --- | --- | --- | --- |
| *Tupaia belangeri* | | | |
| ENaC α | forward reverse | AACGACTGACGACCAACCAA GAAGCTTCTAGGCTGCGGAA | 373 |
| ENaC β | forward reverse | ATTGCTACTCGGATCTGCGG GACAACTCCTTTCCGGCTCA | 206 |
| ENaC γ | forward reverse | GCCCAGCCAACAGTATCGAG GAAAGTGGGTGGGTCATCGT | 260 |
| ENaC δ | forward reverse | GATCAGTACTGGCGCTACCC CCGGTGCTGTTACACAGTCT | 320 |
| GAPDH | forward reverse | GCCTGGAGAAAGCTGCCAAAT ACTGTCAAGGAGGGGAGCTT | 378 |
| *Gallus gallus* | | | |
| ENaC α | forward reverse | CCGCGGTGGTTCTGTGAAG ACCACCAGAGAGAGGCCATT | 939 |
| ENaC β | forward reverse | GATCCTGGCGTGTCTCTACG CAGGCAAGCCTGGAAAAAGG | 448 |
| ENaC γ | forward reverse | GTGACATCGACAGGAGCCAA CAGAAGGATGACGAGCGTGT | 621 |
| ENaC δ | forward reverse | GAGGAGCCAGGGATACCATC CATCCTTTAAGCCTGTTTCTTCCTG | 424 |
| GAPDH | forward reverse | CACTATCTTCCAGGAGCGTGA TTGGCTGGTTTCTCCAGACG | 540 |
