## Supplementary file 8 for "Varying selection pressure for a Na^+^ sensing site in epithelial Na^+^ channel subunits reflect divergent roles in Na^+^ homeostasis"

Alignment of *Marmosa mexicana* sequences (NCBI BioProject PRJNA1052681) with *Homo sapiens* ENaC subunits [NCBI accession numbers NP_001029.1 (ENaC α), NP_000327.2 (ENaC β), NP_001030.2 (ENaC γ)]. No sequences for the δ subunit were identified from *Marmosa mexicana* kidney tissue.

```
Marmosa_alpha      --------------------MKEEKLEREGQHPEPGGPPE-EEEEEGLIEFHRSYRELFQ
NP_001029.1        MEGNKLEEQDSSPPQSTPGLMKGNKREEQGLGPEPAAPQQPTAEEEALIEFHRSYRELFE
                                       ** :* * :*  ***..* :   ***.************:

Marmosa_alpha      FFCNHTTIHGAIRLVCSKHNRMKTAFWAVLWICTFSMMYWQFALLFGEYFSYPVNLNINL
NP_001029.1        FFCNNTTIHGAIRLVCSQHNRMKTAFWAVLWLCTFGMMYWQFGLLFGEYFSYPVSLNINL
                   ****:************:*************:***.******.***********.*****

Marmosa_alpha      NSDKLVFPAVTVCTLNPYRYTKIQEELEELDRITEKTLFDLYKYNSSRIPSNKPRPRRDL
NP_001029.1        NSDKLVFPAVTICTLNPYRYPEIKEELEELDRITEQTLFDLYKYSSFTTLVAGSRSRRDL
                   ***********:********.:*:***********:********.*       .*.****

Marmosa_alpha      QNTLPYPLLMIQNPQSLHHHR-----ASGVQENNPQVDKNDWKIGFILCNKNKSDCFYQT
NP_001029.1        RGTLPHPLQRLRVPPPPHGARRARSVASSLRDNNPQVDWKDWKIGFQLCNQNKSDCFYQT
                   ..***:**  :. * . *  *     **.:.:******.:****** ***:*********

Marmosa_alpha      YSSGVDAVREWYRFHFINILARLDSQ--DLDEAALGNFIFACRFNQASCNQGNYSQFHHP
NP_001029.1        YSSGVDAVREWYRFHYINILSRLPETLPSLEEDTLGNFIFACRFNQVSCNQANYSHFHHP
                   ***************:****:** .   .*:* :************.****.***:****

Marmosa_alpha      VYGNCYTFNGKNNSNLWMSSTPGINNGLSLTLRTERNDFIPLLSTVTGARVMVHGQDEPP
NP_001029.1        MYGNCYTFNDKNNSNLWMSSMPGINNGLSLMLRAEQNDFIPLLSTVTGARVMVHGQDEPA
                   :********.********** ********* **:*.***********************.

Marmosa_alpha      FMDDGGFNLRPGVETSISMRKETLDRLGGNYGDCTKNGSEIQVENIYSSKYTQQVCIHSC
NP_001029.1        FMDDGGFNLRPGVETSISMRKETLDRLGGDYGDCTKNGSDVPVENLYPSKYTQQVCIHSC
                   *****************************:*********:: ***:*.************

Marmosa_alpha      FQESMIRECGCAYMSYPKRDGVEFCDYKKHTAWGYCYYKLQVAFSSDNLGCFAKCRKPCS
NP_001029.1        FQESMIKECGCAYIFYPRPQNVEYCDYRKHSSWGYCYYKLQVDFSSDHLGCFTKCRKPCS
                   ******.******: **. :.**:***.**::********** ****:****:*******

Marmosa_alpha      VTNYQLSAGYSRWPSATSQDWVFQMLSLQNNYTISSK-SGVAKLNIFFKELNYKANSESP
NP_001029.1        VTSYQLSAGYSRWPSVTSQEWVFQMLSRQNNYTVNNKRNGVAKVNIFFKELNYKTNSESP
                   **.************.***:******* *****:..* .****:**********:*****

Marmosa_alpha      SVTMVTLLSNLGSQWSLWFGSSVLSVVEMAELIFDFLVITFLLLLRRLRSRYWAPGHSAQ
NP_001029.1        SVTMVTLLSNLGSQWSLWFGSSVLSVVEMAELVFDLLVIMFLMLLRRFRSRYWSPGRGGR
                   ********************************:**:*** **:****:*****:**....

Marmosa_alpha      GSQEVA--MESSPPSRFSCHS-GASSDQLGPEPSAPV--PPPAYATLTPESVS-------
NP_001029.1        GAQEVASTLASSPPSHFCPHPMSLSLSQPGPAPSPALTAPPPAYATLGPRPSPGGSAGAS
                   *:****  : *****.*. *. . * .* ** **..:  ******** * . .

Marmosa_alpha      ---------
NP_001029.1        SSTCPLGGP
```

```
Marmosa_beta      MNLKKYLVKCLHRLQKGPGYTYKELLVWYCDNTNTHGPKRIICEGPKK------------
NP_000327.2       MHVKKYLLKGLHRLQKGPGYTYKELLVWYCDNTNTHGPKRIICEGPKKKAMWFLLTLLFA
                  *::****:* **************************************

Marmosa_beta      ------------------------------------------------------------
NP_000327.2       ALVCWQWGIFIRTYLSWEVSVSLSVGFKTMDFPAVTICNASPFKYSKIKHLLKDLDELME


Marmosa_beta      ------------------------------------------------------------
NP_000327.2       AVLERILAPELSHANATRNLNFSIWNHTPLVLIDERNPHHPMVLDLFGDNHNGLTSSSAS


Marmosa_beta      --------------------------------QAMKEWYILQSTSILSQVPLEERVQMGY
NP_000327.2       EKICNAHGCKMAMRLCSLNRTQCTFRNFTSATQALTEWYILQATNIFAQVPQQELVEMSY
                                                  **:.******:*.*::*** :* *:*.*

Marmosa_beta      PADQMILACLFGAEPCNHRNFTAIFHPDYGNCYVFNWGMRGKALPSSNPGTEFGLKLILD
NP_000327.2       PGEQMILACLFGAEPCNYRNFTSIFYPHYGNCYIFNWGMTEKALPSANPGTEFGLKLILD
                  *.:**************:****:**:* *****:*****  *****:*************

Marmosa_beta      IDQQDYVHYLTSTAGVRLMLHEQKAYPFLKDGGIYAMPGTET------------------
NP_000327.2       IGQEDYVPFLASTAGVRLMLHEQRSYPFIRDEGIYAMSGTETSIGVLVDKLQRMGEPYSP
                  *.*:*** :*:************.:***:.* *****.****

Marmosa_beta      --------------------------------------------LYPLPKGEKYCNNQDF
NP_000327.2       CTVNGSEVPVQNFYSDYNTTYSIQACLRSCFQDHMIRNCNCGHYLYPLPRGEKYCNNRDF
                                                              *****.*******.**

Marmosa_beta      PDWAYCYSALRMSVLQRETCINGCKESCNDTQYKMTISMADWPSEASEDWIFHVLSYERD
NP_000327.2       PDWAHCYSDLQMSVAQRETCIGMCKESCNDTQYKMTISMADWPSEASEDWIFHVLSQERD
                  ****:*** *.*** ******. ********************************* ***

Marmosa_beta      KTTNLTIDRKGIVKLNIYFQEFNYR-----------------------------------
NP_000327.2       QSTNITLSRKGIVKLNIYFQEFNYRTIEESAANNIVWLLSNLGGQFGFWMGGSVLCLIEF
                  ::**:*:.*****************

Marmosa_beta      ------------------------------------------------------------
NP_000327.2       GEIIIDFVWITIIKLVALAKSLRQRRAQASYAGPPPTVAELVEAHTNFGFQPDTAPRSPN


Marmosa_beta      ----------------------------------------
NP_000327.2       TGPYPSEQALPIPGTPPPNYDSLRLQPLDVIESDSEGDAI
```

```
Marmosa_gamma      MAPGEKITAKIKKNLVVTGPQAPSIKELMKWYCLNTNTHGCRRIVVSRGRLRRLIWIILT
NP_001030.2        MAPGEKIKAKIKKNLPVTGPQAPTIKELMRWYCLNTNTHGCRRIVVSRGRLRRLLWIGFT
                   *******.******* *******:*****.************************:** :*

Marmosa_gamma      LSAVGLILWQCALLVLSFYTVSVSIKVHFQKLNFPAVTICNINPYKYSAVKELLAGLDQE
NP_001030.2        LTAVALILWQCALLVFSFYTVSVSIKVHFRKLDFPAVTICNINPYKYSTVRHLLADLEQE
                   *:**.**********:*************.**:***************:*. ***.*:**

Marmosa_gamma      TKNALKNLYGLSNIKSR-------------------------------------------
NP_001030.2        TREALKSLYGFPESRKRREAESWNSVSEGKQPRFSHRIPLLIFDQDEKGKARDFFTGRKR
                   *.:***.***:.: ..*

Marmosa_gamma      ---------------------VGFQMCANNGTSNDCATYTFNSGVNAIREWYKLHYMNIM
NP_001030.2        KVGGSIIHKASNVMHIESKQVVGFQLCSND--TSDCATYTFSSGINAIQEWYKLHYMNIM
                                        ****:*:*:  :.*******.**:***.***********

Marmosa_gamma      AQVPLEKKINMSYSAEELLVTCFFDGVSCDARNFTLFHHPMYGNCYTFNGATNETILSTS
NP_001030.2        AQVPLEKKINMSYSAEELLVTCFFDGVSCDARNFTLFHHPMHGNCYTFNNRENETILSTS
                   *****************************************:*******.  ********

Marmosa_gamma      MGGSEYGLQVVLYIDEEEYNPFLISSTGAKVVVHRQDEYPFIEDIGTEIETAMATSIGMH
NP_001030.2        MGGSEYGLQVILYINEEEYNPFLVSSTGAKVIIHRQDEYPFVEDVGTEIETAMVTSIGMH
                   **********:***:********:*******::********:**:********.******

Marmosa_gamma      LTESFKLSEPYSQCTKDGSDVPVKSIYNATYSLQICLHSCFQRHMAEKCQCAQYSQPAPK
NP_001030.2        LTESFKLSEPYSQCTEDGSDVPIRNIYNAAYSLQICLHSCFQTKMVEKCGCAQYSQPLPP
                   ***************:******:..****:************ :*.*** ******* *

Marmosa_gamma      NVSYCNYQKHPNWMYCYYKLHQAFVQEELGCQAICREPCNFKEWSLTTSLAQWPSDISEK
NP_001030.2        AANYCNYQQHPNWMYCYYQLHRAFVQEELGCQSVCKEACSFKEWTLTTSLAQWPSVVSEK
                    ..*****:*********:**.**********::*.*.*.****:********** :***

Marmosa_gamma      WMLDVLTWDKGQKENKRLNKTDLAKLLIFYKDLNR-------------------------
NP_001030.2        WLLPVLTWDQGRQVNKKLNKTDLAKLLIFYKDLNQRSIMESPANSIEMLLSNFGGQLGLW
                   *:* *****:*.: **.*****************.

Marmosa_gamma      --------------------CIIARRFWQKTKKWWAQRKTQGQSSSENPEGRTGCDNPTC
NP_001030.2        MSCSVVCVIEIIEVFFIDFFSIIARRQWQKAKEWWAWK--QAPPCPEAPRSPQGQDNPAL
                                       .***** ***:*:***..  *. ...* * .  * ***:

Marmosa_gamma      VLDDDLPTFNTALQLPQALGGHVPGTPPPKYNTLRIDRAFSNQLEDTQ-PEKP
NP_001030.2        DIDDDLPTFNSALHLPPALGTQVPGTPPPKYNTLRLERAFSNQLTDTQMLDEL
                    :********:**:** *** :*************::******* ***  ::
```
